# Expression of short hairpin RNAs using the compact architecture of retroviral microRNA genes

**DOI:** 10.1101/131581

**Authors:** James M. Burke, Rodney P. Kincaid, Francesca Aloisio, Nicole Welch, Christopher S. Sullivan

**Affiliations:** The University of Texas at Austin, Institute for Cellular and Molecular Biology, Center for Synthetic and Systems Biology, Center for Infectious Disease and Dept. Molecular Biosciences, 1 University Station A5000, Austin TX 78712-0162

**Keywords:** shRNA, RNAi, Adeno-associated virus vector, minicircle DNA, BLV, PARP13, p53, pRb

## Abstract

Short hairpin RNAs (shRNAs) are effective in generating stable repression of gene expression. RNA polymerase III (RNAP III) type III promoters (U6 or H1) are typically used to drive shRNA expression. While useful for some knockdown applications, the robust expression of U6/H1-driven shRNAs can induce toxicity and generate heterogeneous small RNAs with undesirable off-target effects. Additionally, the U6 and H1 promoters encompass the majority of the ∼270 base pairs (bp) of vector space required for shRNA expression. This can limit the efficacy and/or number of delivery vector options, particularly when delivery of multiple gene/shRNA combinations is required. Here, we develop a compact shRNA (cshRNA) expression system based on retroviral microRNA (miRNA) gene architecture that uses RNAP III type II promoters. We demonstrate that cshRNAs coded from as little as 100 bps of total coding space can precisely generate small interfering RNAs (siRNAs) that are active in the RNA-induced silencing complex (RISC). We provide an algorithm with a user-friendly interface to design cshRNAs for desired target genes. This cshRNA expression system reduces the coding space required for shRNA expression by greater than two-fold as compared to the U6/H1 promoters, which may facilitate therapeutic RNAi applications where delivery vector space is limiting.

## Introduction

RNA interference (RNAi) is a post-transcriptional mechanism for regulating gene expression in eukaryotes [for review see (1)]. During RNA interference (RNAi) in mammals, small interfering RNAs (siRNAs; ∼22 nts in length) guide the RNA-induced silencing complex (RISC) to mRNAs via base-pair complementarity. RISC association with mRNAs results in cleavage of the mRNA transcript by Argonaute 2 (Ago2), resulting in repression of gene expression (2). The ability of engineered RNAi to specifically target almost any mRNA transcript is a potentially useful therapeutic agent for a broad spectrum of human diseases, including cancer, inflammatory diseases, neurodegenerative diseases, ocular diseases, and viral infections (3-6). However, strategies for successful implementation of therapeutic RNAi in humans continue to be refined.

To achieve RNAi, small double-stranded RNAs (∼22 nts in length) need to be delivered to or generated in the cytosol of the target cell. This can be accomplished via transfection of synthetic siRNAs into cells, or by transfection or transduction of DNA templates that express siRNA-generating RNAs (6). One advantage of siRNA generation from a DNA template [e.g., lentiviruses, plasmids, Adeno-associated virus (AAV) vectors, PCR amplicons, and small circular DNAs (coligos)] is the potential for stable long-term RNAi (6,7). A common method to achieve this is through expression of short hairpin RNAs (shRNAs), which are processed by the RNase III enzyme Dicer into siRNAs. Typically, the U6 or H1 promoters/enhancers (RNAP III class III) are used to promote RNAP III-mediated transcription of the shRNA (8,9). These promoters are located entirely upstream of the transcription initiation site and promote robust shRNA expression. This makes design of the downstream shRNA straightforward and modular, while ensuring consistent and robust shRNA expression. However, the upstream U6/H1 promoters (215, 246 bps respectively) encompass >80% of the genomic template space required for expression of a shRNA. Additionally, the robust shRNA expression promoted by the U6/H1 promoters can result in cytotoxicity by saturating factors required for generating endogenous microRNAs (miRNAs). Furthermore, these shRNA vectors often give rise to abundant small RNAs with heterogeneous 5’ ends that can increase undesirable off-target effects on unintended mRNA targets (10,11). Thus, alternative shRNA expression strategies may offer advantages over the conventional U6/H1-driven shRNA for *in vivo* applications whereby smaller templates, less robust expression, and more precise biogenesis are important.

Bovine leukemia virus (BLV) expresses five pre-miRNAs from ∼550 bps of genomic space (12). Each pre-miRNA is directly transcribed by RNAP III from individual compact RNAP III type II genes (13-15). This is analogous to typical shRNA generation, but unlike the U6/H1 promoters, the two *cis* promoter elements (the A and B boxes) that promote RNAP III transcription initiation are located within or directly downstream of the pre-miRNA hairpin (13,16,17). This architecture requires as little as 80-bps of template space to encode for the pre-miRNA and promoter elements. Further, BLV pre-miRNA expression appears to be less robust in comparison to U6/H1-driven shRNAs, which may account for the lack of noticeable cytotoxicity accompanied with BLV pre-miRNA expression (12,13). This attribute of BLV pre-miRNA expression is likely important, since cytotoxicity during BLV persistence *in vivo* would be disadvantageous to viral fitness.

Here, we develop an shRNA expression system based on the design principles gleaned from the architecture of the BLV miRNA genes. We demonstrate the efficacy of these BLV-based compact shRNA cassettes (cshRNAs) by designing multiple cshRNAs and evaluating their expression and RNAi activity against three different target transcripts. This work provides a method to express shRNA-generated siRNAs with defined termini from a coding sequence of 100 bps, which may be useful for RNAi-based therapies and laboratory applications where smaller gene silencing expression cassettes are desirable.

## Materials and Methods

### Plasmids

The pBLV-B1 vector is described in (13). To generate the 115-bp and 100-bp BLV-B1 templates, the 115-bp BLV-B1 sense primer or 100-bp BLV-B1 sense primer was used with the BLV-B1 antisense primer (Supplementary Table S1) to generate the amplicons from the pBLV-B1 via PCR using Phusion polymerase (New England BioLabs). Amplicons were digested with Xho1/Xba1 and ligated into the *Xho1/Xba1* sites of the pIDTSmart-kan (pISK), in which we previously engineered *Xho1/Xba1* sites (12). Each cshRNA construct was generated by annealing sense and antisense oligonucleotides (Integrated DNA technologies; Supplementary Table S1) that encoded the cshRNA hairpin and ∼15 nts of flanking sequence. Phusion polymerase was used to fill-in the single-stranded overhangs. The cshRNA-encoding fragments were then extended via PCR using the universal B1-flanking sense and antisense primers (Supplementary Table S1) to add the ∼40 bps of the BLV-B1 locus flanking the BLV-pre-miR-B1 hairpin and the *Xho1* and *Xba1* sites. The amplicons were then digested with Xho1/Xba1 and ligated into *Xho1/Xba1* sites of the pIDT-smart-Kan (pISK). The RISC reports were made by extension of RISC reporter (RR) primers (Supplementary Table S1) with Phusion polymerase. Amplicons were digested with Xho1/Xba1 and ligated into the *Xho1/Xba1* sites of pcDNA3.1dsRluc (18). For the p53-and pRb-3’-UTR reporters, the p53-and pRb-3’UTRs were amplified using Phusion polymerase and the p53-and Rb-3’UTR primers (Supplementary Table S1) from cDNA prepared from HEK293T RNA using Superscript III reverse transcriptase (Invitrogen) and polyT_20_ primer. The amplicons were digested with Spe1/Sal1 and ligated into the *Xho1/Xba1* sites of pcDNA3.1dsRluc (Addgene plasmid # 68053). All plasmids were sequence verified by ICMB core facilities (The University of Texas at Austin).

### Cell culture

HEK293T and HEK293 cells were obtained from ATCC. Cells were maintained in DMEM supplemented with FBS (Cellgro; 10% vol/vol). Transfections were performed using Lipofectamine 2000 (Invitrogen) according to manufacturer’s instructions.

### cshRNA target prediction algorithm

A web based graphical user interface for the identification of target sites and generation of complete cshRNA designs is available at: http://w3.biosci.utexas.edu/sullivanlab/cshRNA. Source code is available at: https://github.com/rpkincaid/cshRNA.

### Northern blotting

HEK293T cells (6-well format; ∼70% confluent) were transfected with 1-ug of cshRNA expression vectors using Lipofectamine 2000 (Invitrogen). Forty-eight hours post transfection, RNA was isolated using PIG-B (19). Northern blot analysis was performed as described in (20,21). Briefly, 15-ug of RNA was fractionated on 15% PAGE-UREA gel and transferred to Amersham Hybond –N+ membrane (GE Healthcare). The membrane was then probed in ExpressHyb hybridization solution (Clontech, CA) with a pool of indicated DNA oligos (Integrated DNA technologies; Supplemental Table 1) radio-labeled with [γ-32P]-ATP (6000 Ci/mmol) by T4 polynucleotide kinase (New England Biolabs). Membranes were exposed to a storage phosphor screen (GE Healthcare) and visualized with a Typhoon biomolecular imager (GE Healthcare). Membranes were stripped by incubation in boiling 0.1% SDS with agitation for five minutes.

### RISC reporter assays

HEK293T cells (24-well format, 70% confluent) were co-transfected with 500-ng/well of either pIDTSmart-kan (pISK-EV) or each cshRNA vector and 2.5-ng/well of the firefly luciferase expression vector (pcDNA3.1dsLuc2CP) and 2.5-ng/well of each individual Rluc RISC reporter using 2-ul of lipofectamine 2000/well. Forty-eight hours post-transfection, lysates were harvested and dual luciferase assay was performed on a Luminoskan Ascent Luminometer (Thermo Electronic). Raw Renilla/firefly ratios were normalized to pISK-EV and Rluc vector 3’-UTR for each data set. For each experimental replicate, transfections were performed in triplicate.

### Immunoblot analysis

HEK293 cells (12-well format; 60% confluent) were transfected with 1-ug of either pISK (EV) or pISK-cshRNA-p53-h8 vectors using lipofectamine 2000. Forty-eight hours post-transfection, cells were lysed in SDS solution (1% SDS, 2% β-mercaptoethanol) by boiling for ten minutes followed by one minute of vortexing (22). Equal volumes were fractionated on 7.5% or 12.5% PAGE-SDS followed by transfer to a 0.45 um nitrocellulose membrane (Bio Rad). The membrane was probed with p53 monoclonal mouse antibody (IMGENEX: catalog #: GSC-1010) used at 1:1000 dilution, ZC3HAV1 (PARP13) rabbit polyclonal antibody (Gene Tex; catalog #: GTX120134) used at 1:5000 dilution, and α-Tubulin monoclonal mouse antibody (Sigma-Aldrich: catalog #: T6199) used at 1:10,000 dilution. Antibodies were diluted in Phosphate Buffered Saline containing 0.1% Tween 20 (PBST) and 5% BSA. After 3 washes with PBST, membranes were blotted with IRDye 800CW and IRDye 680LT secondary antibodies (LI-COR) diluted 1:10,000 in PBST with 5% BSA. Membranes were washed four times with PBST and then scanned on an Odyssey CLx infrared imaging system (LI-COR).

### RNA sequencing

HEK293T cells in a 6-well format were transfected with 1-ug of an individual cshRNA vector. Forty-eight hours post-transfection, small RNAs (<70 nts) were isolated from total RNA as described in (20) and pooled. Small RNA libraries were prepared for Illumina small RNA sequencing from the pooled small RNAs using the multiplex small RNA library prep set (New England Bio Labs) and sequenced on a Illumina HiSeq 2500 and NextSeq 500 platforms (GSAF, UT Austin). Adapter sequences were trimmed from the reads using fastx clipper from the FASTX-Toolkit software (http://hannonlab.cshl.edu/fastx_toolkit). The preprocessed reads were then mapped to cshRNA reference sequences using SHRiMP2 software package (23). Small RNA reads were quantitated using custom scripts.

### Structural prediction

Secondary structure predictions of cshRNAs were generated using the RNA folding form on the Mfold web server (24).

### Results and Discussion

#### Design principles of an shRNA template based on BLV miRNA genes

We set out to develop a smaller shRNA expression system by imitating the design principles of the BLV miRNA gene architecture (Figure 1A). To engineer shRNAs that utilize similar features of the BLV pre-miRNAs, we chose BLV-pre-miR-B1 as the archetype because it is readily transcribed by RNAP III, is efficiently processed by Dicer, and generates a highly dominant 3p derivative (Figure 1B) (12,13). We used the position of the 3p arm of BLV-pre-miR-B1 to serve as the siRNA template. To allow for variation of the siRNA template while still maintaining efficient transcription and shRNA structure, we restricted the siRNA sequences to those whose sequence complement (permitting G-U wobble pairing) conserves the minimal consensus nucleotides within A-box-1 on the 5p arm (nucleotides: 1,2,9,10,11; Figure 1C). We also conserved siRNA nucleotides 21-23 (TCT), which are part of the termination signal, to ensure proper transcription termination to generate the 2-nt overhang. We permitted variation of the remaining nucleotides in the siRNA. These parameters give rise to permissible variable siRNAs with the following sequences: 5’-YYNNNNNNYYRNNNNNNNNNUCU-3’ (Y = C/U, R = A/G, N = A/U/G/C).

**Figure 1.**
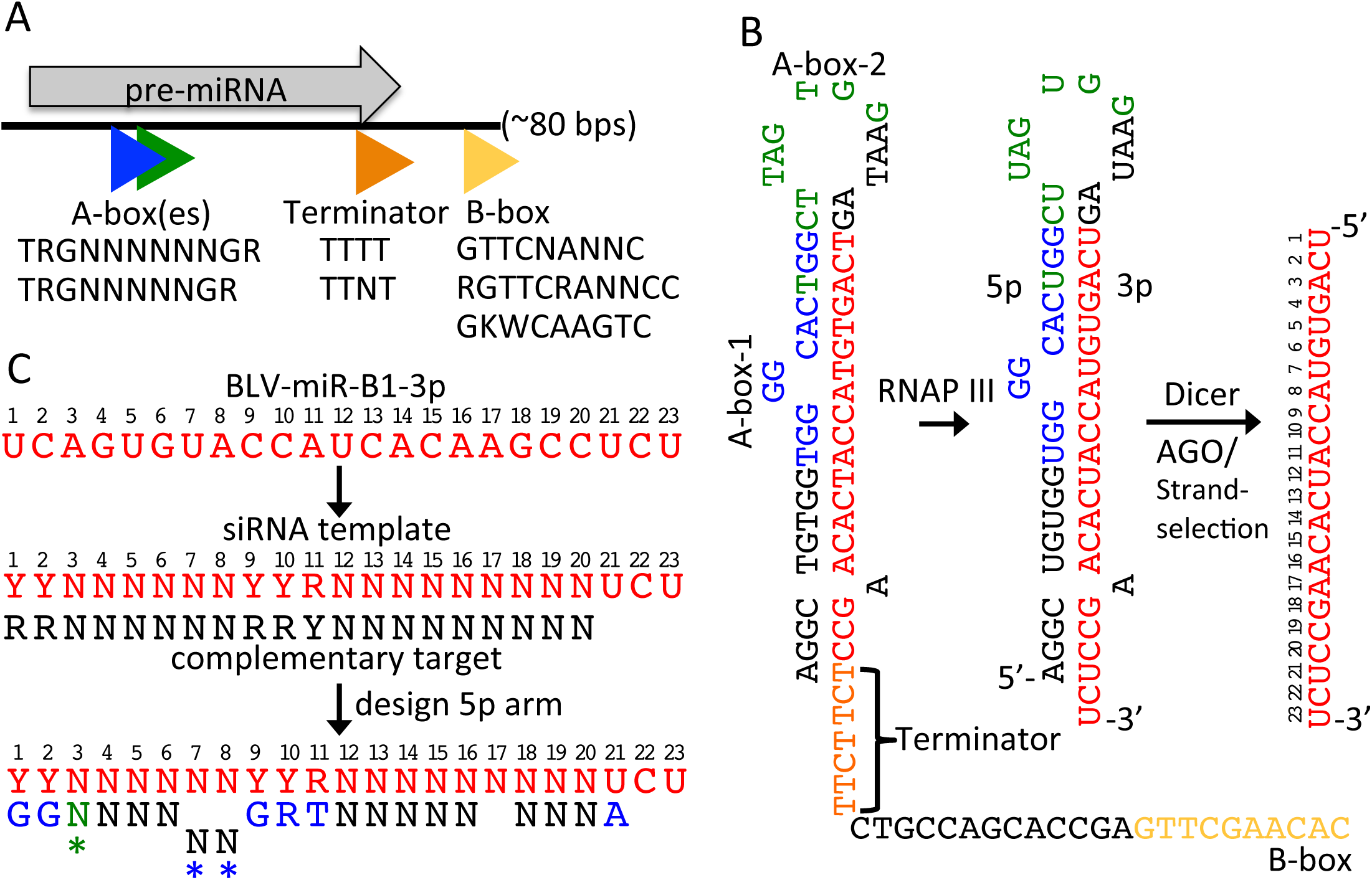
Design of the BLV-base compact shRNA system. (A) Schematic of BLV miRNA gene architecture with the consensus sequences for each RNAP III type II promoter element shown relative to the pre-miRNA hairpin. (B) Illustration of the positions of the RNAP III transcriptional elements [A-box-1 (blue), A-box-2 (green), terminator (orange)] relative to the BLV-miR-B1 precursor. The nucleotides of BLV-miR-B1-3p are shown in red. (C) The design strategy for the compact shRNA (cshRNA) expression system. The 3p arm of BLV-pre-miR-B1 serves as the siRNA template. To conserve shRNA structure and the consensus nucleotides in A-box-1, positions 1, 2, 9, 10, 11 of the siRNA were restricted to maintain base pairing (including G-U wobble base pairing) with the consensus nucleotides of the A-box-1 element in the 5p arm. To maintain the position of transcription termination, nucleotides 21, 22, 23 of BLV-miR-B1-3p were conserved in the siRNA template. These allow for permissible siRNAs that conform to the template: 5’-YYNNNNNNYYRNNNNNNNNNUCU-3’. Sequences that were perfectly complementary to the first 20 nts of the siRNA template were then identified in a transcript of interest. Once a compatible target sequence with the siRNA template was identified in a transcript, the siRNA and the corresponding 5p arm containing the consensus nucleotides of A-box-1 were designed. Asterisks indicate variations in the design used to generate cshRNAs described in the following sections. The green asterisk and nucleotide indicates that the consensus T of A-box-2 was conserved in some cshRNAs, which restricts position 3 of the siRNA to a purine to maintain base pairing in these cshRNAs. The blue asterisks indicate that mismatching the nucleotides at positions 7 and 8, as in BLV-pre-miR-B1, was maintained in some cshRNAs.

To test the efficacy of this design, we searched for sequences perfectly complementary to nucleotides 1-20 of the siRNA template (5’-NNNNNNNNNYRRNNNNNNRR-3’) in three different sequences: the 3’ UTR of *RB1,* the 3’UTR of *TP53* and the entire mRNA of *ZC3HAV1* (*PARP13)*. In order increase the number of possible cshRNA targets/transcript, we did not require complementarity between nucleotides 21-23 of the siRNA and the target mRNA. We note that base pairing between the 3’-end of siRNAs and target mRNAs is not required for mRNA recognition, translational repression, or Ago2 slicer activity (25,26). Using these parameters, numerous sites complementary to the siRNA template were identified in each transcript (Supplementary Table S2), demonstrating that target sequences amenable to our cshRNA design constraints frequently occur in diverse target sequences. Permissible target sequences in each transcript were assigned arbitrary identification numbers (e.g. cshRNA-target-h1). We selected 27 total cshRNA targets (8 targeting the *p53* 3’UTR, 4 targeting the *RB* 3’UTR, and 15 targeting the *PARP13* mRNA) for analysis.

To make the cshRNA-expressing cassettes, we selected complementary siRNAs and designed the corresponding 5p arms containing the consensus nucleotides of the A-box elements (Figure 1C). For some constructs (the p53-and RB-targeting cshRNAs), we preserved the mismatched nucleotides at positions 7 and 8 of BLV-pre-miR-B1 that result in a predicted symmetrical “bulge” in the stem of shRNA hairpin. However, for the PARP13-targeting cshRNAs, this mismatch design element was omitted. In all cshRNAs, we maintained the predicted single asymmetric nucleotide bulge of the BLV-B1 pre-miRNA hairpin by not allowing a complementary nucleotide for position 17 of the siRNA (Figure 1C). For all constructs we also conserved the A in the 5p arm that is complementary to the U at siRNA position 21. The rationale for this was two-fold: 1) It preserves the 2-nt 3’ overhang of the shRNA, and 2) It provides the purine in the pyrimidine/purine junction (-1/+1) that guides the position of RNAP III transcription initiation at the 5’-end of the shRNA. The siRNA and corresponding 5p arm were then inserted into the BLV-pre-miR-B1 hairpin in place of the BLV-pre-miR 5p and 3p arms. The predicted secondary structure of the cshRNAs we chose for analysis largely maintained the structure of BLV-pre-miR-B1 (Figure 2A-D), indicating that Dicer-mediated processing of the shRNAs would likely be preserved.

**Figure 2.**
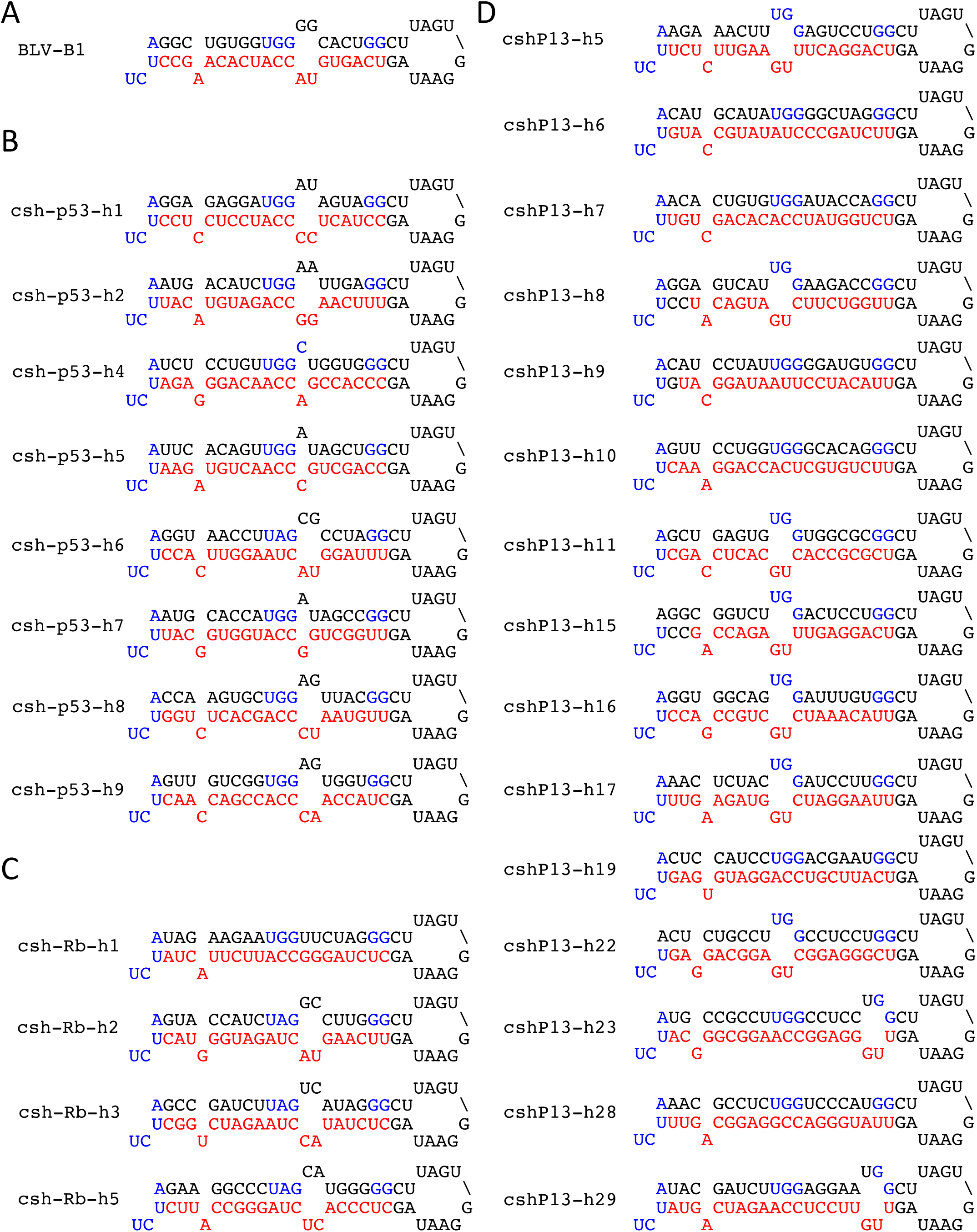
Structural predication of cshRNAs. Secondary structure predications of (A) BLV-pre-miR-B1, (B) *p53*-targeting cshRNAs, (C) *RB1*-targeting cshRNAs, (D) *PARP13*-targeting cshRNAs. BLV-miR-B1-3p and the siRNAs are shown in red. The nucleotides of the RNAP III transcriptional elements (+1 purine, A-box-1, terminator signal) are in blue.

#### BLV-based shRNA templates generate specifically processed and RISC-active siRNAs

To assess cshRNA expression and biogenesis, we made cshRNA expression vectors by replacing the BLV-pre-miR-B1 hairpin with each cshRNA in the pBLV-B1 vector, which contains 130 bps of the BLV-miR-B1 locus in a plasmid that lacks a known mammalian promoter (pIDTSmart-kan) (13). These cassettes included the cshRNA (56 bps), 43 bps upstream of BLV-pre-miR-B1, and 31 bps downstream of BLV-pre-miR-B1. We transfected HEK293T cells with each individual cshRNA vector. Northern blot analysis revealed that most (26/27) of the cshRNAs express RNAs consistent with the size of the shRNA hairpin, and the majority (23/27) generate a readily detectable derivative small RNA consistent with the size of a siRNA (∼22 nts; Figure 3A-C). These data demonstrate that our cshRNA design strategy is effective at generating shRNAs that produce small RNAs with diverse sequences.

**Figure 3.**
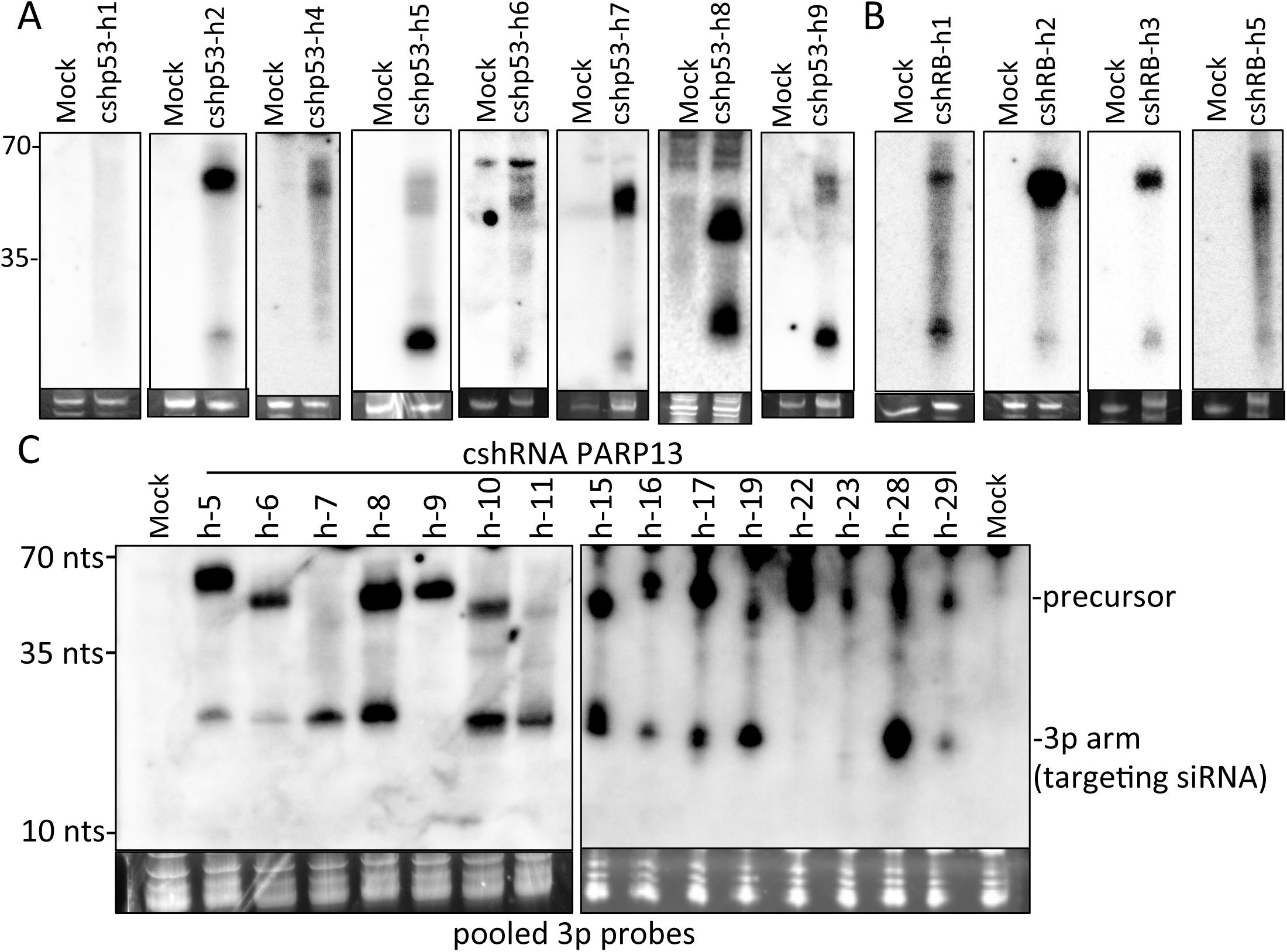
Northern blot analysis of p53-, pRb-, and PARP13-targeting cshRNAs. Northern blot analysis of the (A) p53-, (B) pRb-, and (C) PARP13-targeting cshRNAs. HEK293T cells were transfected with indicated cshRNA vectors. Forty-eight hours later, RNA was isolated and subject to northern blot analysis. The membrane was blotted with pooled probes complementary to the 3p arms of each cshRNA. Mock lanes indicate cells that were not transfected with a cshRNA expression vector. Below the northern blots are small RNAs stained with ethidium bromide (EtBr) to serve as a loading control.

We performed Illumina high-throughput small RNA sequencing (RNA-seq) to precisely determine the sequence of the small RNA derivatives from our cshRNA constructs. All the *p53*-targeting cshRNAs generated small RNAs that map to the 5p and/or 3p arms of their respective cshRNA. The majority (5/8) produced dominant 3p siRNAs (Figure 4A). CshRNA-p53-h5 produced co-dominant 5p and 3p derivatives. CshRNAs p53-h1 and p53-h4 produced dominant 5p derivatives, likely accounting for the lack of observable 3p siRNAs by northern blot analysis (Figure 3A). Importantly, the most abundant siRNA isoform for approximately half of the constructs had the intended single predominant 5’ and 3’ termini (Figure 4A). Three of the four *RB*-targeting cshRNAs produced dominant 3p siRNAs, while cshRNA-RB-h1 produced co-dominant 5p and 3p derivatives (Figure 4B). All the *RB*-targeting cshRNAs generated siRNAs with the correct 5’-end for the most abundant isoform. We did not obtain small RNA reads mapping to cshRNA-PARP13-h9, which is consistent with our northern blot analysis (Figure 3C). However, cshRNA-PARP13-h9 does express an shRNA precursor (Figure 3C). This suggests that Dicer may not efficiently process cshRNA-PARP13-h9 precursor RNA into siRNAs. All the other *PARP13*-targeting cshRNA constructs (14/15) generated small RNAs that map to the 5p and/or 3p arms of the pre-miRNA (Figure 4C). *PARP13*-targeting cshRNAs h5, h6, h7, h8, h10, h11, h16, h17, h19, and h28 all produced dominant 3p siRNAs, whereas cshRNAs h15, h19, h22, h23, and h29 generated co-dominant levels of 3p and 5p arms. Small RNA reads mapping to *PARP13*-targeting cshRNAs 22 and 23 were low, likely accounting for the lack of observable small RNAs by northern blot analysis. We observed abundant total reads of the 3p arms generated from *PARP13*-targeting cshRNAs h5, h8, h10, h15, h16, h17, h19, and h28, suggesting these cshRNAs are efficiently transcribed, processed, and loaded into Argonaute. Importantly, the most abundant 3p arm isoform generated from the majority of *PARP13*-targeting cshRNAs had the correct intended 5’ and 3’ termini. These data indicate that most cshRNAs generate siRNAs of the intended sequence, a desirable feature for minimizing off-targeting effects.

**Figure 4.**
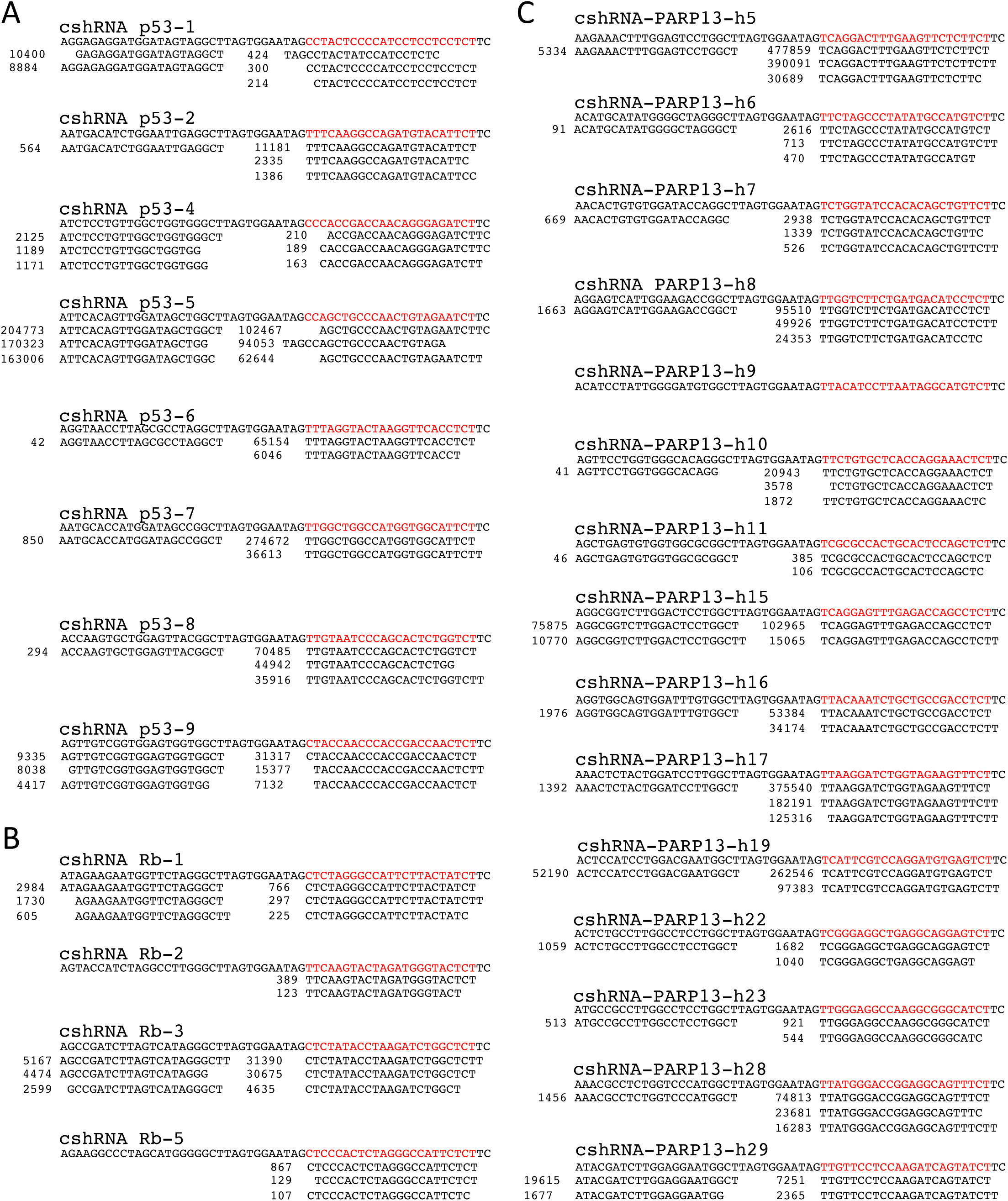
Small RNA sequencing of cshRNA-generated small RNAs. Small RNA sequencing of small RNAs generated from cshRNAs specific to (A) p53, (B) pRb, (C) PARP13 transcripts. HEK293T cells were transfected with individual cshRNA vectors. Forty-eight hours later, small RNAs were isolated from cells and pooled for small RNA sequencing library preparation. Small RNAs were sequenced on the Illumina HiSeq 2500 and NextSeq 500 platforms. Shown is the reference sequence for each cshRNA with the nucleotides of the intended siRNA in red. Below each reference sequence are the total read counts of the most abundant small RNAs that align to the 5p and 3p arms of the indicated cshRNAs.

We next assayed whether cshRNA-generated small RNAs are active in RISC. We engineered luciferase-based RISC reporters for the cshRNA-pRb-3, cshRNA-p53-8, and cshRNA-p53-9 by inserting two perfectly complementary sequences to either the 5p or 3p arms of each cshRNA into the 3’-UTR of *Renilla* luciferase (Figure 5A). For the PARP13-targeting cshRNAs, we engineered similar reporters for the siRNA (3p arm). Co-expression of cshRNA-Rb-h3 resulted in a significant reduction (∼80%) of luciferase activity of the 3p reporter, but had no effect on the 5p reporter (Figure 5B). Similar results were observed for cshRNA-p53-h8 (Figure 5C). CshRNA-p53-h9 repressed both the 5p and 3p reporters (Figure 5D), indicating that cshRNA-p53-h9 produces co-dominant siRNAs, consistent with the abundant read counts of both the 5p and 3p derivatives in our RNA-seq analysis (Figure 4). The majority (11/14) of *PARP13*-targeting cshRNAs reduced Rluc expression from their respective RISC reporters by greater than 50%, with six decreasing Rluc expression by over 80% (Figure 5E). These data demonstrate that cshRNAs generate RISC-active siRNAs.

**Figure 5.**
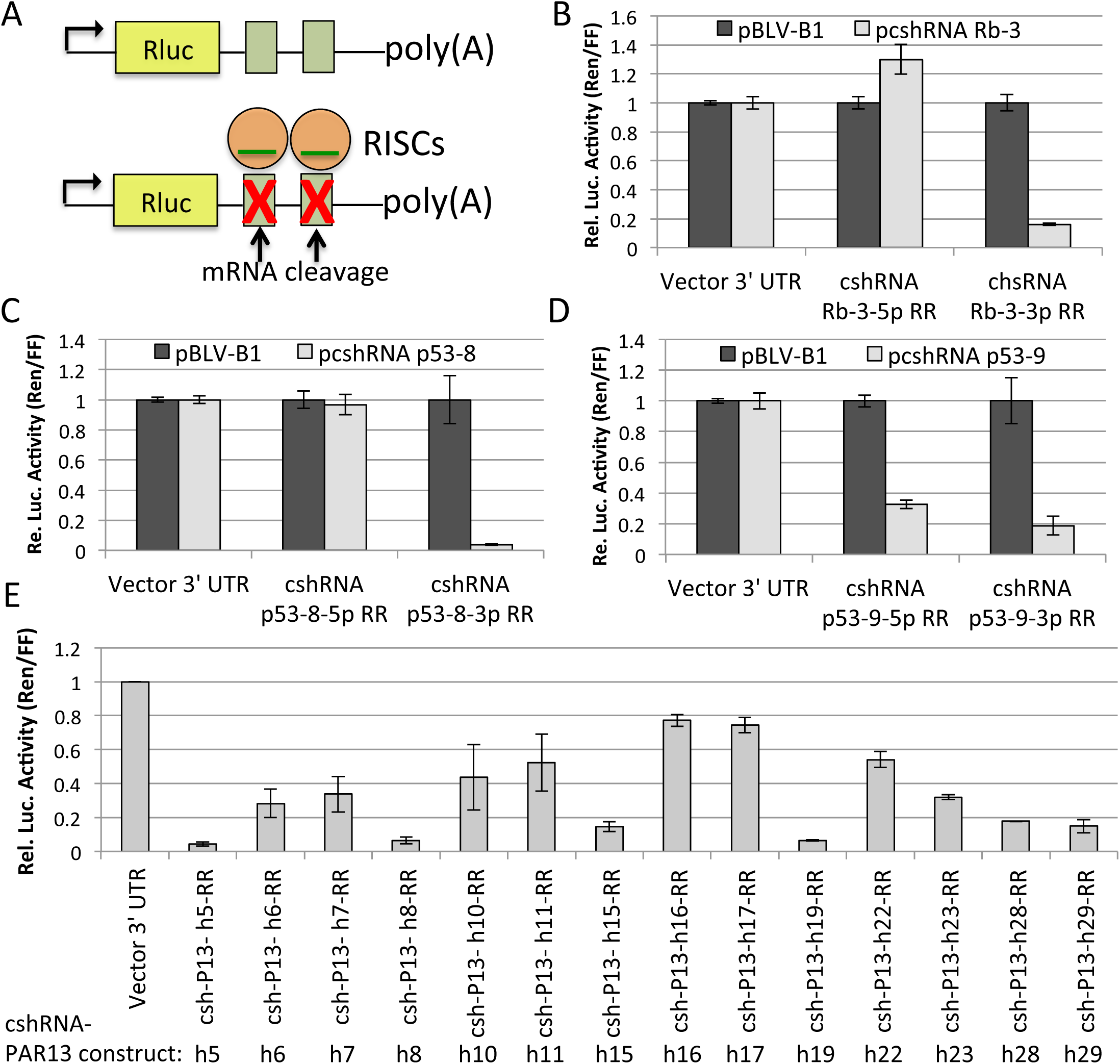
Analysis of cshRNA-generated small RNA RISC activity. (A) Diagram of the RISC reporter cleavage assay. Green boxes represent complementary sequences to siRNAs. (B) Luciferase assay to measure RISC activity of the 5p and 3p arms of the cshRNA-Rb-3. HEK293T cells were co-transfected with the indicated vectors in triplicate, and luciferase assays were performed forty-eight hours post-transfection. The graph represents the average +/-the SD. (C and D) Similar to (B) with the indicated cshRNA vectors and respective RISC reporters. (E) RISC reporter assay to test RISC activity of the PARP13-targeting cshRNAs. HEK293T cells were co-transfected with either negative control vector (pIDTSmart-kan) or each individual PARP13-targeting cshRNA vector and either negative control reporter (vector 3’-UTR) or the respective cshRNA RISC reporter, as indicated on graph. The bars represent the average +/-SD of three independent experiments, in which the transfections were performed in triplicate in each experiment. The data was normalized to pIDTSmart-kan (EV; data not shown) and the vector 3’ UTR.

To assay whether cshRNA-generated siRNAs can repress their intended target within the context of a longer natural 3’ UTR, we inserted the entire 3’ UTRs of the *p53* and *RB* mRNAs downstream of *Renilla luciferase* in pcDNA3.1dsRluc and evaluated the ability of the corresponding cshRNAs to repress Rluc expression (Figure 6A). All four of the *RB*-targeting cshRNAs decreased Rluc-RB-3’-UTR expression, with cshRNA-RB-h3 reducing Rluc-pRB-3’-UTR expression to the largest degree (80% reduction; Figure 6B). Roughly half of the *p53*-targeting cshRNAs (h6, h7, h8, and h9) repressed Rluc-p53-3’-UTR, with cshRNA-p53-h8 reducing expression by 80% (Figure 6C). To determine if cshRNAs are able to knockdown endogenous target proteins, we evaluated p53 and PARP13 levels in cells transiently expressing either cshRNA-p53-h8 or any of several different *PARP13*-targeting cshRNAs (h7, h8, h11, h19). Both cshRNA-p53-h8 and cshRNA-PARP13-h19 significantly reduced p53 and PARP13 levels, respectively (Figure 6D,E). Combined, these results demonstrate that cshRNAs can knockdown gene expression of their intended target mRNA transcript.

**Figure 6.**
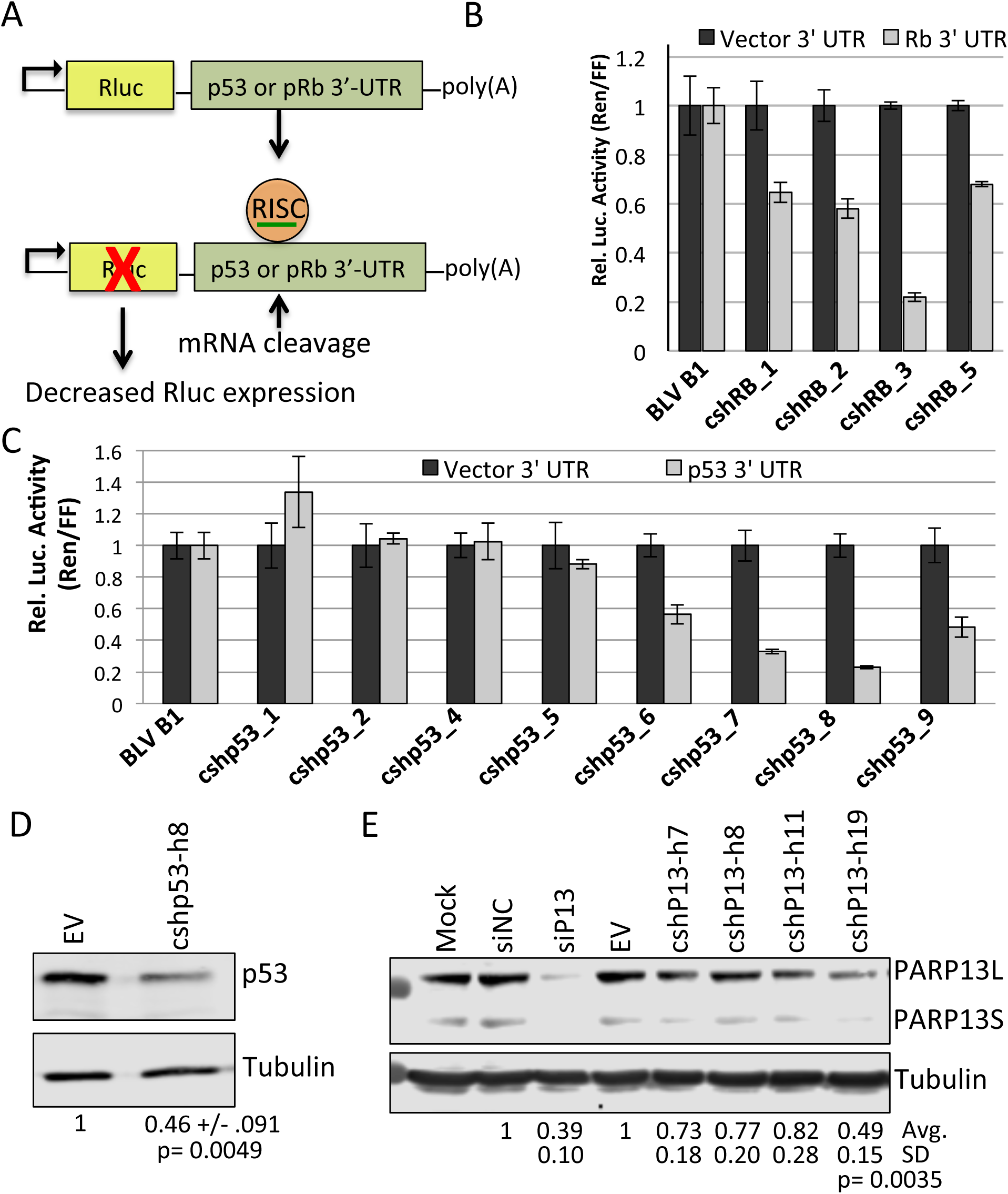
cshRNA-generated small RNAs can downregulate targets. (A) Schematic of luciferase reporter assay to assess cshRNA-mediated repression of the p53 and pRb 3’ UTR. (B and C) Luciferase assays to measure the ability of the (B) *RB*-targeting cshRNAs and (C) *p53*-targeting cshRNAs to target the 3’-UTRs of the *RB* and the *p53* mRNAs. The bars represent the average luciferase ratio (Ren/FF) +/-the SD of representative experiments performed in triplicate. (D and E) Immunoblot analysis to assay knockdown of endogenous p53 (D) or PARP13 (E). HEK293 cells were transfected with 1-ug of a negative control vector pIDTSmart-kan (EV) or the indicated cshRNA vectors. 48 hours post-transfection, equal volumes were subjected to immunoblot analysis. The average fold-decrease +/-SD in p53 (n=3) or PARP13 (n=4) relative to tubulin is shown below the blots. For unknown reasons, the positive control PARP13 siRNAs did not reduce PARP13 levels in some experiments. The results from these experiments were excluded from these analyses. The p-values were calculated using one sample t-test.

### The cshRNA coding space can be reduced to 100 base pairs

To determine if we could reduce the template space required for cshRNA expression, we tested whether we could eliminate sequence from the 5’ flanking region of BLV-miR-B1 locus or select cshRNAs. Although some RNAP III type II promoters contain a functional upstream TATA box at approximately position -30 (16,17), the BLV-miR-B1 locus and our cshRNA vectors lack this *cis* element, indicating the upstream sequence may not be important for RNAP III transcription initiation. Therefore, we made vectors in which we truncated 15-or 30-bps from the 5’-end of the upstream of BLV-pre-miR-B1/cshRNA in the pBLV-B1, cshRNA-p53-h8, and cshRNA-PARP13-h19 vectors (Figure 7A). Northern blot analysis revealed that deletions of this region did not markedly reduce expression of BLV-miR-B1 or the cshRNA-generated siRNAs (Figure 7B,C). Thus, small RNAs can be efficiently expressed from a 100 bp gene expression cassette that includes the 56-bp cshRNA, 31 bps downstream, and only 13 bp of upstream BLV-pre-miR-B1 sequence.

**Figure 7.**
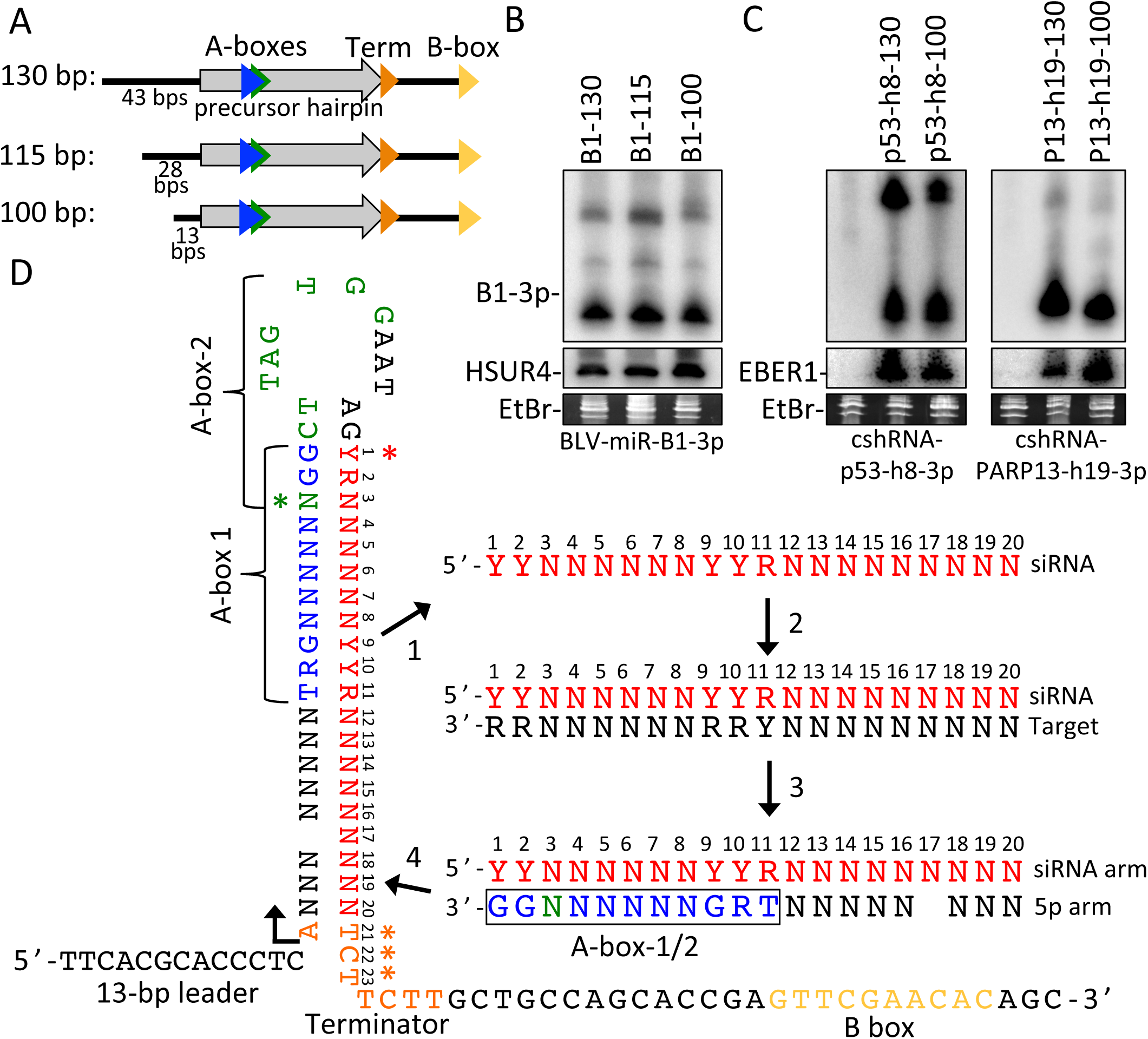
Optimized cshRNA design. (A) Illustration of RNAP III transcriptional elements in BLV-miR-B1 locus. The 43-bp sequence upstream of BLV-pre-miR-B1 in pBLV-B1 (130-bp template) was truncated by 15 bps or 30 bps to generate 115-or 100-bp templates in the pBLV-B1, cshRNA-P53-h8, and cshRNA-PARP13-h19 expression vectors. (B) Northern blot analysis of RNA harvested from HEK293T co-transfected with the 130-bp, 115-bp, and 100-bp BLV-miR-B1 templates in the pIDTSmart-kan vector and the pHSUR4 transfection control. (C) Northern blot analysis of RNA harvested from HEK293T co-transfected with the 130-bp and 100-bp cshRNA-p53-h8 and cshRNA-PARP13-h19 templates in the pIDTSmart-kan vector and the EBER1 transfection control. (D) Illustration of the cshRNA design template. The siRNA template allows for sequence variation while maintain base pairing (including G-U wobble) with the A box elements in the 5p arm (step 1). Perfectly complementary sequences to the first 20 nts of the siRNA template are identified in a target mRNA (step 2). Once a compatible target is identified, the siRNA and corresponding 5p arm, which includes conservation of consensus nucleotides of A-box-1 and removal of the nucleotide that would base pair with position 17 of the siRNA, are designed (step 3). The siRNA and corresponding 5p arm are then inserted into the BLV-miR-B1 backbone template, which includes the 13 bp 5’ leader, the A at the transcription start site, the terminator signal, and the B box promoter element (step 4). The asterisks indicate nucleotides that can be optimized to increase siRNA biogenesis and/or RISC activity. The red asterisk indicates that small RNAs with a 5’-U at position 1 of the siRNAs may promote more efficient biogenesis of the siRNAs, but is not absolutely required. The green asterisk shows that the consensus T of A-box-2 may promote more efficient and consistent expression of cshRNAs, but is not required. The orange asterisks indicate the nucleotides in terminator signal that are in the siRNA but that we did not require to base pair with the target mRNA in the default setting.

### Algorithm for easy and flexible design of cshRNA templates

We generated a computational design tool for cshRNA expression cassettes. The web-based interface program is available at http://w3.biosci.utexas.edu/sullivanlab/cshRNA. Source code is available at https://github.com/rpkincaid/cshRNA. The default setting generates the 100-bp template shown in Figure 7D, but there is an optional parameter that can incorporate the additional 30-bp sequence upstream of BLV-pre-miR-B1 (add 5’ flanking sequence). Although not supported by the vectors we tested (Figure 7A-C), its plausible that additional BLV 5’ cis elements may improve expression of some cshRNAs. To simplify the cshRNA design, we omitted the requirement to mismatch nucleotides at positions 7 and 8 (Figure 7D), as this was not required for the biogenesis of the *PARP13*-targeting cshRNAs (Figures 2-4). The default setting maximizes the number of possible cshRNA targets/transcript by using the least stringent yet still permissible siRNA template. However, the program includes several optional design parameters that restrict the siRNA template to potentially optimize cshRNA effectiveness. These optional parameters include: i) Conservation of the consensus thymidine in A-box-2 [complementary to siRNA position 3 (Figure 7D)], which may provide more robust and/or consistent shRNA expression (13), though it was not absolutely required for cshRNA expression (e.g. cshRNA-p53-h8). ii) The requirement for the siRNA to contain a 5’-uracil. This appeared to promote more efficient and more precise siRNA biogenesis (Figure 3), which is consistent with studies demonstrating a 5’-U promotes Dicer processing and Argonaute association (27,28). iii) The requirement for nucleotides 21-23 of the siRNA to be perfectly complementary to the target mRNA. Although not absolutely required for cshRNA-mediated knockdown of gene expression [e.g. shRNA-p53-h8 and cshRNA-PARP13-h19 (Figure 6D,E)], this may increase knockdown efficiency due to more efficient Ago2 slicer activity in some contexts (25,26). We note that selecting this option substantially decreases the number of possible cshRNA targets/transcript. Finally, we included the option of excluding cshRNAs with internal termination signal-like sequences and repetitive sequences, which are likely problematic to effective cshRNA generation. Combined, these features provide flexibility in cshRNA design that allows for modulation of cshRNA biogenesis/effectiveness while maximizing potential target sequences.

## Conclusions

Most of the cshRNA cassettes we tested generated RISC-active siRNAs (Figure 3-6). However, a small fraction of the cshRNAs did not generate abundant siRNAs. It is likely that variability in the A-box promoter elements and variability in shRNA structure and processing account for this. Therefore, for routine laboratory purposes where template size is not an issue, U6/H1-driven shRNAs may be preferable to cshRNAs. In addition, not all RISC-active cshRNA-generated siRNAs reduced gene expression of their target mRNA (Figure 6). This is likely a consequence of inaccessibility of the mRNA target sequence to RISC. Therefore, cshRNA effectiveness could be enhanced by combining our design program with existing siRNA target accessibility prediction programs (29).

Reducing the coding space required for small RNA expression may be especially useful for therapeutic applications. Recently, Mefferd et al. (2015) used the biogenesis mechanisms of the mouse γ-herpesvirus 68 (MHV68) miRNAs to develop CRISPR/Cas9 single guide RNAs (sgRNAs) expressed by tRNA promoters (RNAP III type II) to facilitate AAV-based Cas9/sgRNA vectors (30-35). In a similar manner, our BLV-based cshRNA expression system uses RNAP III type II promoters to drive shRNA expression. This requires less than half of the space of U6/H1-driven shRNA cassettes. We envision that cshRNAs may be useful for small payload gene therapy vectors that express multiple shRNAs or combinations of protein coding genes and shRNAs. Such vectors include supercoiled minicircle DNA vectors, which can be only a few hundred nucleotides of total size, and AAV vectors which have less than 5 kilobases of payload capacity (36-44). In summary, our BLV-based cshRNA template reduces the amount of coding sequence required for shRNA expression, and may offer other benefits in terms of amount of expression and specific shRNA processing. cshRNAs show promise as a tool for future therapeutic RNAi applications.

## Funding

This work was supported by a Burroughs Wellcome Investigators in Pathogenesis Award; and a grant from the Cancer Prevention and Research Institute of Texas [RP140842].

## Acknowledgements

The authors would like to thank Dr. Joan Steitz for the pEBV RIJ plasmid.

